# Reward-Based Equilibrium Propagation

**DOI:** 10.64898/2026.07.30.741817

**Authors:** Yoshimasa Kubo

**Affiliations:** Department of Computer Science, Lakehead University, Thunder Bay, Canada

## Abstract

Equilibrium propagation (EP) is a biologically plausible alternative to backpropagation for training neural networks. EP typically relies on free and nudged dynamical phases: during the free phase, the network relaxes toward an equilibrium state, whereas during nudging, the output state is perturbed toward a target using a teaching signal. However, it remains unclear whether the brain has access to such explicit target signals. Inspired by Attention-Gated Brain Propagation (BrainProp), a reward-based learning framework proposed by Pozzi et al. (2020), we introduce a reward-based variant of EP that replaces full target-based nudging with a selected-output binary reward signal. The proposed method updates the network using only the chosen class and whether that choice is correct, without directly revealing the full target vector. We evaluate the method on MNIST, Fashion-MNIST, and CIFAR-10 using both multilayer perceptrons and convolutional neural networks. The proposed reward-based EP achieves performance close to that of conventional EP across all three datasets, although it generally converges more slowly during the early stages of training. Generalization-gap analyses show similar behavior for the two methods on MNIST and Fashion-MNIST, while reward-based EP exhibits a smaller training–test accuracy gap during later training on CIFAR-10. We further investigate the effect of the exploration probability used during stochastic class selection and find that moderate exploration can provide small performance improvements, although its effect is dataset-dependent. These results demonstrate that EP can learn effectively from sparse, action-specific reward feedback rather than a complete supervised target.

## 1 Introduction

Equilibrium Propagation (EP) [Scellier and Bengio, 2017, 2019, Ernoult et al., 2019, Laborieux et al., 2021, Laborieux and Zenke, 2022] is a biologically plausible alternative to backpropagation (BP) for training neural networks. EP has attracted considerable attention because it enables synaptic updates based on locally available quantities while achieving competitive performance on a range of supervised learning tasks. In particular, recent advances have shown that convolutional neural networks trained with improved variants of EP can attain performance comparable to that of networks trained with BP [Laborieux et al., 2021].

EP typically relies on free and nudged dynamical phases. During the free phase, the network evolves in the absence of an external teaching signal until its internal states approach an equilibrium. During nudging, a small perturbation is introduced at the output layer through a task-dependent loss function, commonly the squared-error loss, so that the network state is driven toward a desired target. Synaptic weights are then updated based on differences between the corresponding neural activities. Because these updates depend on locally available pre- and postsynaptic quantities, EP is often described as implementing a local Hebbian-like learning mechanism.

Despite this biological motivation, conventional EP still assumes access to explicit output targets. It remains unclear how such target signals would be represented or generated in the brain [Lee et al., 2014]. In particular, it is not obvious how a biologically realistic system could construct a supervised loss from a complete target vector and use it to perturb the activities of all output neurons.

A related line of work has explored reward-based learning mechanisms that avoid direct access to explicit output targets. Attention-Gated Brain Propagation (BrainProp) [Pozzi et al., 2020] is one such framework. Instead of receiving the correct target vector, the network selects an output and receives a scalar reward indicating whether the selected response was correct. BrainProp combines this global reward signal with feedback connections that propagate action-specific credit-assignment signals from higher to lower processing levels. These feedback signals preferentially target the neurons and synapses that contributed to the selected action, thereby gating plasticity within the relevant pathways [Roelfsema and Holtmaat, 2018, Roelfsema et al., 1998, Pooresmaeili et al., 2014]. This framework is biologically motivated by interactions between neuromodulatory reward signals, such as those associated with dopaminergic systems, and top-down attentional feedback that regulates synaptic plasticity [Schultz et al., 1997, Schultz, 2002, Friedrich et al., 2010, Huang et al., 2013].

In this study, we introduce reward-based Equilibrium Propagation (REP) by adapting the selected-output reward formulation of BrainProp to the symmetric three-phase EP framework. Rather than providing the network with a complete target vector, the network selects a class and receives a binary reward indicating whether the selected class is correct. A reward-based loss is defined only for the selected output unit, which is nudged according to the discrepancy between its activity and the received reward. The resulting action-specific perturbation propagates through the recurrent dynamics of EP, enabling credit assignment to lower layers without explicitly revealing the correct class to the network.

We evaluate REP on MNIST, Fashion-MNIST, and CIFAR-10 using both multilayer perceptrons and convolutional neural networks, and compare its performance with that of conventional target-based EP. In addition to maximum test accuracy and learning dynamics, we analyze the gap between training and test accuracy throughout training. This analysis allows us to examine whether replacing the complete target vector with an action-specific reward signal changes the relationship between fitting the training data and performance on unseen examples. We also investigate how the exploration probability used during stochastic class selection influences learning performance and variability across datasets.

Our results show that REP achieves performance close to that of conventional EP across all three datasets, despite receiving substantially less supervisory information. REP generally converges more slowly during the early stages of training, but it approaches the performance of EP on MNIST and Fashion-MNIST. The two methods exhibit similar generalization gaps on these datasets, whereas REP maintains a smaller training–test accuracy gap during later training on CIFAR-10. These findings demonstrate that symmetric three-phase EP can learn from sparse, action-specific binary reward feedback rather than a complete supervised target vector.

## 2 Methods

### 2.1 Equilibrium Propagation

Equilibrium Propagation (EP) is a learning algorithm for convergent recurrent neural networks in which neural states evolve toward steady states through recurrent dynamics [Scellier and Bengio, 2017, 2019, Ernoult et al., 2019]. Let **x** denote an input, **s** the collection of neural states, and ***θ*** the trainable parameters. The network dynamics are defined through a primitive function Φ(**x, s, *θ***). In a discrete-time formulation, the neural states are updated according to

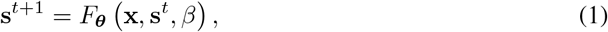

where *β* controls the strength and direction of an external nudging signal.

In conventional supervised EP, the task-dependent cost is defined using the output state **y** and a complete target vector **t**. A commonly used choice is the squared-error cost

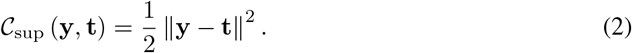

The negative gradient of this cost with respect to the output state is

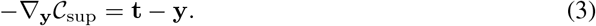

Thus, conventional supervised EP nudges the output layer using the errors of all output units and therefore requires access to the complete target vector.

Standard EP begins with a free phase. During this phase, the nudging factor is set to zero, *β* = 0, and the network evolves for *T*_free_ time steps:

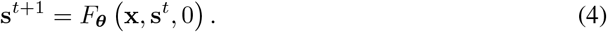

Because no external cost is applied, the network evolves only according to its internal dynamics. The resulting steady state is denoted by

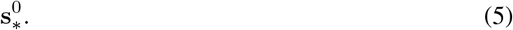

The output component of this state determines the network prediction before any teaching signal is introduced.

In conventional two-phase EP, the free phase is followed by a single nudged phase in which a small nonzero value of *β* weakly perturbs the output state toward the target. The difference between the free and nudged steady states is then used to estimate the task gradient. However, this two-phase estimator is affected by a first-order bias when a finite nudging strength is used.

To reduce this bias, Laborieux et al. [2021] introduced a symmetric three-phase formulation that uses both positive and negative nudging. After the free phase, the network is first evolved from the free steady state using a positive nudging factor +*β*:

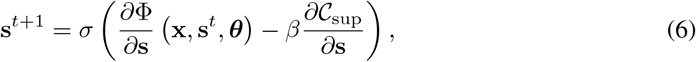

where *σ*(·) denotes the neuronal activation function. The perturbation term

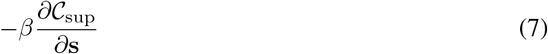

drives the network toward a state with a lower supervised cost. After *T*_nudged_ time steps, the network reaches the positively nudged steady state

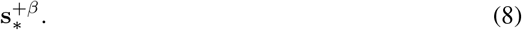

The network is then evolved from the free steady state using the opposite nudging factor −*β*:

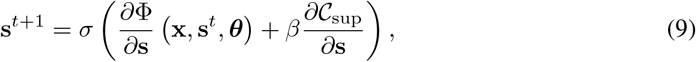

producing the negatively nudged steady state

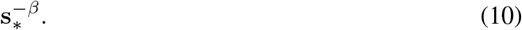

The same input **x** and target **t** are used in both nudged phases, while only the sign of the nudging factor is reversed.

The symmetric EP parameter update is then computed as

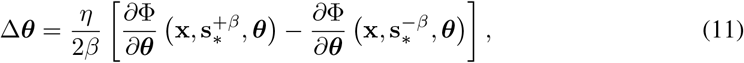

where *η* is the learning rate. Using nudging factors with opposite signs cancels the leading finite-*β* bias and provides a more accurate estimate of the gradient [Laborieux et al., 2021].

For a fully connected weight matrix **W**^*l*^ connecting layers *l* and *l* + 1, the corresponding local update can be written as

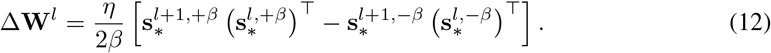

This contrastive update depends only on the activities of neurons connected by the corresponding synapses in the two nudged phases. In this study, we use this symmetric three-phase formulation for both conventional supervised EP and the proposed reward-based EP. The distinction between the two methods lies in the cost function used in Equations 6 and 9: conventional EP uses the complete target-based cost in Equation 2, whereas the proposed method replaces it with a selected-output reward cost.

### 2.2 Attention-Gated Brain Propagation

Attention-Gated Brain Propagation (BrainProp) is a reward-based learning framework that trains neural networks without revealing a complete target vector [Pozzi et al., 2020]. For an input **x**, the network computes an output activity vector

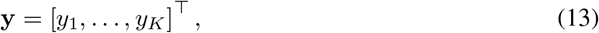

where each output unit corresponds to a possible class or action. The network then selects a class *a* using a stochastic action-selection mechanism.

BrainProp employs a Max–Boltzmann controller. With probability 1 − *ϵ*, the output unit with the highest activity is selected. With probability *ϵ*, a class is sampled from a categorical distribution whose probabilities are obtained by applying the softmax function to the output activities:

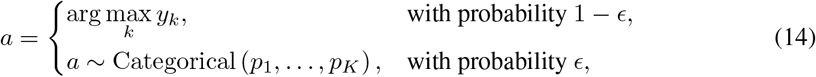

where

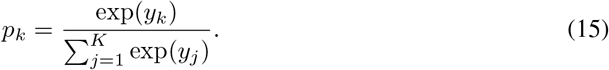

The original BrainProp study used *ϵ* = 0.02, corresponding to greedy selection in 98% of trials and Boltzmann sampling in the remaining 2% [Pozzi et al., 2020]. We use the same action-selection formulation, with the output activities serving directly as the logits of the categorical distribution.

After a class has been selected, the network receives a binary reward

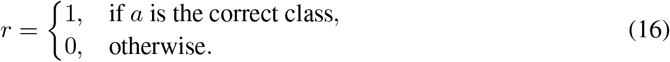

When an incorrect class is selected, the correct class is not revealed to the network. Learning therefore relies only on the selected class and the received reward.

The activity *y*_*a*_ of the selected output unit is interpreted as the expected reward associated with selecting class *a*. The discrepancy between the received and expected reward is

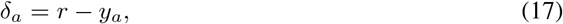

which corresponds to a reward-prediction error. Equivalently, BrainProp defines a squared-error objective only for the selected output unit:

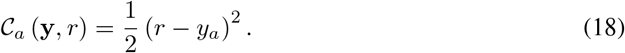

The negative gradient of this objective with respect to the complete output vector is

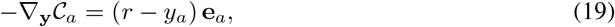

where **e**_*a*_ is the one-hot basis vector associated with the selected class. Consequently, the direct reward-based signal is nonzero only for the selected output unit, while all nonselected output units receive zero direct perturbation.

In the original BrainProp framework, feedback connections explicitly propagate an action-specific attentional signal from the selected output toward lower network layers. This feedback signal identifies the neurons and synapses that contributed to the selected action and thereby gates plasticity within the relevant pathways [Roelfsema and Holtmaat, 2018, Roelfsema et al., 1998, Pooresmaeili et al., 2014]. BrainProp therefore combines a globally available reward-prediction signal with action-specific feedback activity to provide credit assignment without revealing the complete target vector.

In the proposed method, we do not implement BrainProp’s explicit attentional-feedback recursion or its synaptic update rule. Instead, we adapt its stochastic class-selection mechanism and selected-output reward objective to the symmetric three-phase EP framework. In conventional supervised EP,the positive nudged dynamics are defined by Equation 6, while the negative nudged dynamics are defined by Equation 9. Both equations use the complete target-based cost C_sup_ defined in Equation 2.

Consequently, conventional EP requires the complete target vector **t** and directly perturbs all output units according to their target errors.

To obtain the proposed reward-based formulation, we replace the supervised cost *C*_sup_ in both Equation 6 and Equation 9 with the selected-output reward cost *C*_*a*_ defined in Equation 18. Thus, the positive reward-based nudged dynamics are

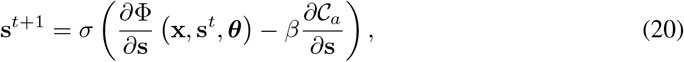

which replace the supervised cost term in Equation 6. The network then evolves toward the positively nudged steady state 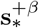.

Similarly, the negative reward-based nudged dynamics are

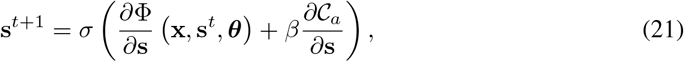

which replace the supervised cost term in Equation 9. The network then evolves toward the negatively nudged steady state 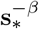.Therefore, the only modification to the nudged dynamics of symmetric EP is the replacement

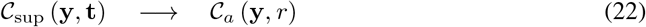

in both the positive and negative nudged phases. The signs of the nudging terms remain unchanged from Equations 6 and 9.

Because *C*_*a*_ depends only on the activity *y*_*a*_ of the selected output unit, the direct nudging signal in both phases is restricted to that unit. During the positive nudged phase, the selected output is perturbed in the direction of the negative cost gradient given in Equation 19. During the negative nudged phase, the sign of this perturbation is reversed. In both phases, the resulting action-specific perturbation propagates to the hidden layers through the recurrent dynamics of the EP network.

The selected class *a* and the corresponding reward *r* are determined from the free-phase output state 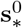 and are held fixed during both nudged phases. Thus, the same selected class and reward are used to compute *C*_*a*_ in Equations 20 and 21; only the sign of the nudging term differs. After the network reaches 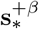 and 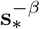, the parameters are updated using the standard symmetric three-phase EP rule in Equation 11.

Thus, BrainProp provides the stochastic class-selection mechanism and selected-output reward formulation, whereas EP provides the recurrent propagation of the output perturbation and the local contrastive parameter-update rule.

### 2.3 Model and Dataset Specifications

We evaluated the proposed reward-based Equilibrium Propagation (REP) method on MNIST, Fashion-MNIST, and CIFAR-10. MNIST and Fashion-MNIST were used to evaluate REP with fully connected multilayer perceptrons, whereas CIFAR-10 was used to examine its applicability to a convolutional architecture. The dataset-specific architectures and training hyperparameters are summarized in Table 1.

**Table 1:** Dataset-specific network architectures and training hyperparameters.

| Hyperparameter | MNIST | Fashion-MNIST | CIFAR-10 |
| --- | --- | --- | --- |
| Network architecture | 784-512-10 | 784-512-256-10 | Conv(64)-Conv(128)-Conv(128)-FC(10) |
| Kernel sizes | — | — | 3, 3, 3 |
| Strides | — | — | 1, 1, 1 |
| Paddings | — | — | 1, 1, 0 |
| Pooling | — | — | Max pooling after each convolutional layer |
| Free-phase iterations, $T_{\text{free}}$ | 60 | 120 | 120 |
| Nudged-phase iterations, $T_{\text{nudged}}$ | 12 | 12 | 12 |
| Mini-batch size | 64 | 64 | 128 |
| Number of epochs | 30 | 30 | 120 |
| Layer-wise learning rates | 0.4, 0.05 | 0.2, 0.08, 0.02 | 0.25, 0.12, 0.08, 0.01 |
| Weight decay | None | None | $3 \times 10^{-4}$ for all layers |
| Learning-rate decay | No | No | Yes |

For MNIST, we used a fully connected network with 784 input units, one hidden layer containing 512 units, and 10 output units. For Fashion-MNIST, we used a deeper fully connected network with two hidden layers containing 512 and 256 units, respectively. For CIFAR-10, we used a convolutional network consisting of three convolutional layers with 64, 128, and 128 channels, respectively, followed by a fully connected output layer with 10 units. Max pooling was applied after each convolutional layer.

All experiments used the symmetric three-phase EP formulation, hard-sigmoid activation functions, and stochastic gradient descent without momentum. The positive and negative nudged phases used nudging factors of +*β* and −*β*, respectively, with |*β*| = 0.1.

For REP, the Max–Boltzmann exploration probability was set to *ϵ* = 0.02, unless otherwise specified. The class selected from the free-phase output and the corresponding binary reward were used to construct the selected-output squared-error cost defined in Equation 18. The selected class and reward were held fixed during both the positive and negative nudged phases.

For conventional supervised EP, hereafter referred to as EP, the complete target vector was used to construct the full target-based squared-error cost defined in Equation 2. Reward-based class selection and Max–Boltzmann exploration were not used for EP.

For each dataset, REP and EP used identical network architectures, phase durations, learning rates, mini-batch sizes, numbers of training epochs, and optimization settings. Thus, the two methods differed only in the cost function and output-level learning signal used during the positive and negative nudged phases.

We report the maximum test accuracy obtained over the fixed training period, following the evaluation convention commonly used in prior EP studies. For each experimental condition, the reported value is the mean of the maximum test accuracies obtained across five random seeds, together with the corresponding standard deviation.

The source code used for all experiments is publicly available at https://github.com/ykubo82/reward_based_ep.

## 3 Results

### 3.1 MNIST

The first row of Table 2 summarizes the results on MNIST. REP achieved a maximum test accuracy of 98.05%, compared with 98.26% for EP. Although EP obtained a slightly higher accuracy, the difference between the two methods was only 0.21 percentage points, indicating that REP remained highly competitive despite using only the selected class and binary reward rather than the complete target vector.

**Table 2:**
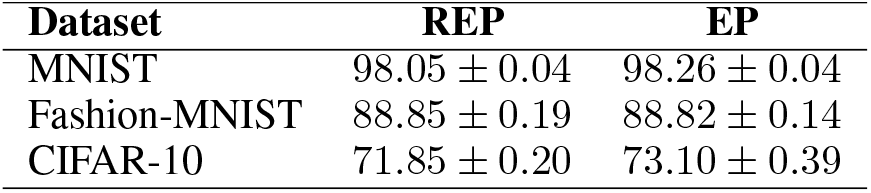
Maximum test accuracy obtained by REP and EP. Results are reported as the mean ± standard deviation over five random seeds. MNIST and Fashion-MNIST models were trained for 30 epochs, whereas the CIFAR-10 model was trained for 120 epochs.

| Dataset | REP | EP |
| --- | --- | --- |
| MNIST | $98.05 \pm 0.04$ | $98.26 \pm 0.04$ |
| Fashion-MNIST | $88.85 \pm 0.19$ | $88.82 \pm 0.14$ |
| CIFAR-10 | $71.85 \pm 0.20$ | $73.10 \pm 0.39$ |

The corresponding test-accuracy learning curves are shown in Figure 1a. EP converged more rapidly during the early epochs, whereas REP exhibited slower initial convergence. However, REP largely closed the performance gap after approximately 10 epochs and subsequently reached a level of test accuracy similar to that of EP.

**Figure 1:**
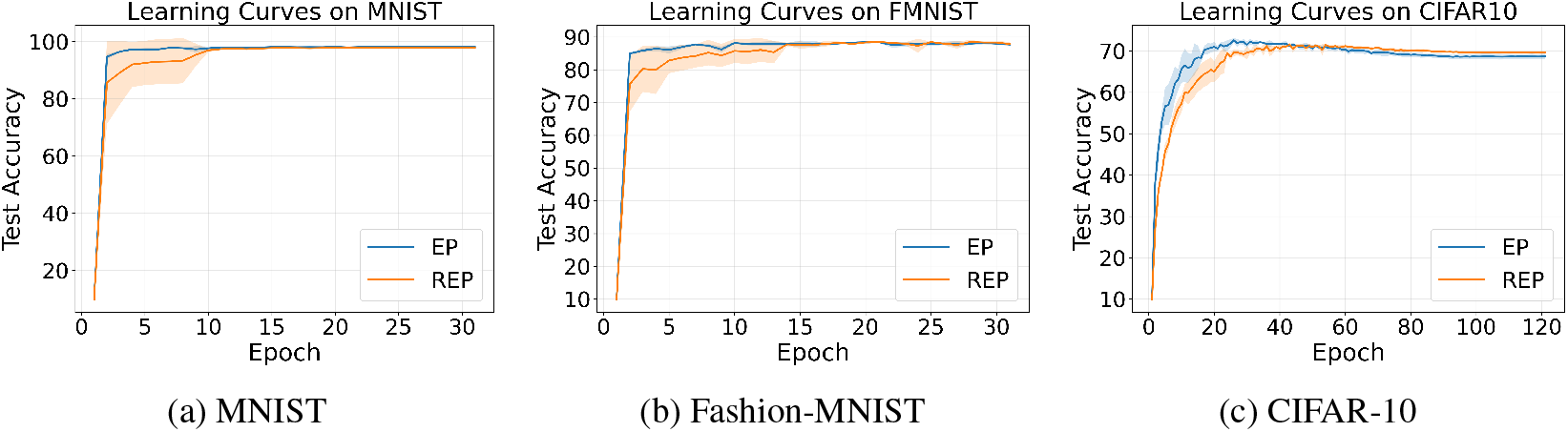
Test-accuracy learning curves for EP and REP on MNIST, Fashion-MNIST, and CIFAR-10. Solid lines indicate the mean test accuracy over five random seeds, and shaded regions indicate the corresponding standard deviation. EP converges more rapidly during the early stages of training on all three datasets. On CIFAR-10, EP test accuracy gradually declines after reaching its peak, whereas REP remains comparatively stable during later epochs.

Figure 2a shows the corresponding generalization gaps, defined as the difference between training and test accuracy at each epoch. REP exhibited greater variability during the early stages of training, whereas the gaps of EP and REP became similar after approximately 10 epochs. During the later stages of training, both methods maintained relatively small and comparable generalization gaps.

**Figure 2:**
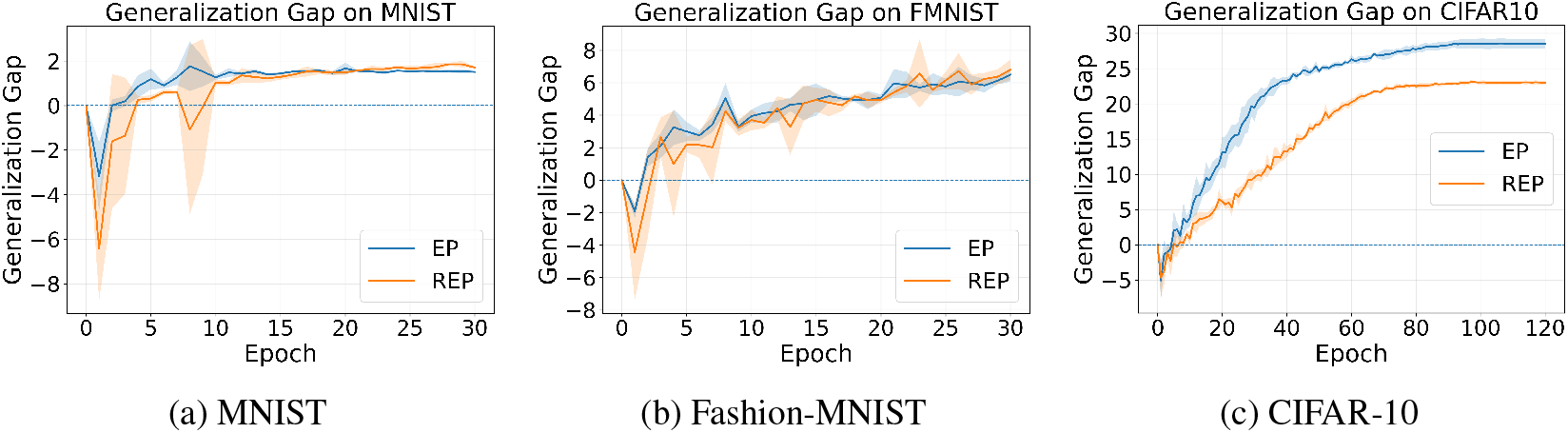
Generalization gaps for EP and REP on MNIST, Fashion-MNIST, and CIFAR-10. The generalization gap is defined as training accuracy minus test accuracy. Solid lines indicate the mean over five random seeds, and shaded regions indicate the corresponding standard deviation.

### 3.2 Fashion-MNIST

The second row of Table 2 presents the results on Fashion-MNIST. REP achieved a maximum test accuracy of 88.85%, while EP achieved 88.82%. The difference between the two methods was therefore negligible, showing that REP can attain performance comparable to conventional EP on a more challenging image-classification task.

Figure 1b shows the corresponding test-accuracy learning curves. As on MNIST, EP converged more rapidly during the early epochs, whereas REP exhibited slower initial convergence. After approximately 15 epochs, however, REP reached a level of test accuracy comparable to that of EP, and the two methods exhibited similar performance during the remainder of training.

The generalization-gap curves in Figure 2b show a similar overall pattern for the two methods. REP exhibited somewhat greater variability during several early and intermediate epochs, but the gaps of EP and REP remained broadly comparable as training progressed. By the end of training, both methods showed similar differences between training and test accuracy.

### 3.3 CIFAR-10

The third row of Table 2 reports the results on CIFAR-10. REP achieved a maximum test accuracy of 71.85%, compared with 73.10% for EP, corresponding to a difference of 1.25 percentage points. Although REP did not match the maximum accuracy of EP, its performance remained reasonably close, demonstrating that the proposed reward-based nudging mechanism can also be applied successfully to a convolutional EP architecture.

The test-accuracy learning curves in Figure 1c reveal a notable difference between the two methods. EP converged more rapidly during the early epochs and reached a higher peak accuracy. After approximately 40 epochs, however, its test accuracy gradually declined. REP also exhibited a small decrease after reaching its peak, but the decline was less pronounced, and its performance remained comparatively stable during the later epochs. Consequently, although EP achieved the higher maximum accuracy, REP retained slightly higher test accuracy near the end of training.

A clearer difference between the two methods is visible in the generalization-gap curves shown in Figure 2c. After approximately 40 epochs, REP consistently exhibited a smaller gap between training and test accuracy than EP. By the end of training, the gap was approximately 28–29 percentage points for EP and approximately 23 percentage points for REP. This result is consistent with the comparatively stable test accuracy of REP during later training. However, the smaller gap may partly reflect lower training accuracy rather than improved generalization alone and should therefore be interpreted cautiously.

### 3.4 Effect of Exploration Probability

In the preceding experiments, the exploration probability was set to *ϵ* = 0.02, following the setting used in the original BrainProp study [Pozzi et al., 2020]. Although that study briefly discussed the role of exploration, it did not systematically examine how different values of *ϵ* affect classification performance. We therefore conducted an ablation study to evaluate the sensitivity of REP to this hyperparameter.

Table 3 reports the maximum test accuracy obtained on each dataset for exploration probabilities ranging from *ϵ* = 0.00 to *ϵ* = 0.05. All other model architectures and training hyperparameters were held fixed. The setting *ϵ* = 0.02, used in the main experiments, is included as the reference condition.

**Table 3:** Exploration ablation results for REP on MNIST, Fashion-MNIST, and CIFAR-10. The exploration probability *ϵ* determines the probability of sampling a class from the Boltzmann distribution rather than selecting the output unit with the highest activity. Results are reported as the maximum test accuracy (%) over training, averaged over five random seeds, with the corresponding standard deviation.

| Exploration probability | MNIST | Fashion-MNIST | CIFAR-10 |
| --- | --- | --- | --- |
| $\epsilon = 0.00$ | $97.85 \pm 0.07$ | $88.71 \pm 0.16$ | $71.84 \pm 0.24$ |
| $\epsilon = 0.01$ | $98.03 \pm 0.11$ | $84.99 \pm 7.58$ | $71.84 \pm 0.25$ |
| $\epsilon = 0.02$ | $98.05 \pm 0.04$ | $88.85 \pm 0.19$ | $71.85 \pm 0.20$ |
| $\epsilon = 0.03$ | $98.09 \pm 0.05$ | $86.91 \pm 3.71$ | $71.72 \pm 0.11$ |
| $\epsilon = 0.04$ | $98.12 \pm 0.05$ | $88.97 \pm 0.15$ | $71.60 \pm 0.35$ |
| $\epsilon = 0.05$ | $98.04 \pm 0.06$ | $88.79 \pm 0.18$ | $72.01 \pm 0.28$ |

On MNIST, fully greedy class selection with *ϵ* = 0.00 produced the lowest mean accuracy of 97.85%. Introducing exploration improved performance across all nonzero values, with the highest mean accuracy of 98.12% obtained at *ϵ* = 0.04. However, the differences among the nonzero exploration settings were relatively small.

A similar overall pattern was observed on Fashion-MNIST. The highest mean accuracy, 88.97%, was obtained at *ϵ* = 0.04, compared with 88.71% under fully greedy selection. Nevertheless, the results at *ϵ* = 0.01 and *ϵ* = 0.03 showed substantially larger variability across random seeds, indicating that some exploration settings may lead to unstable training on this dataset.

On CIFAR-10, the effect of *ϵ* was comparatively limited. Test accuracy remained within a narrow range across the examined values, from 71.60% to 72.01%. The highest mean accuracy was obtained at *ϵ* = 0.05, whereas fully greedy selection achieved 71.84%. These results suggest that moderate exploration can provide a small improvement over fully greedy class selection, although the optimal value of *ϵ* appears to depend on the dataset.

## 4 Discussion

In this study, we introduced reward-based Equilibrium Propagation (REP) by adapting the selected-output reward formulation of BrainProp [Pozzi et al., 2020] to the symmetric three-phase EP framework. Unlike conventional EP, which uses the complete target vector to generate the output perturbation, REP selects a class and receives only a binary reward indicating whether that selection is correct. The resulting reward-based cost is applied only to the selected output unit, and the corresponding perturbation is propagated through the recurrent dynamics of EP.

Across MNIST, Fashion-MNIST, and CIFAR-10, REP achieved performance that was competitive with conventional EP. On MNIST and Fashion-MNIST, the differences in maximum test accuracy between the two methods were small, whereas a larger gap was observed on CIFAR-10. The test-accuracy learning curves further showed that REP generally converged more slowly during the early stages of training. Nevertheless, REP gradually approached the performance of EP on MNIST and Fashion-MNIST and remained comparatively stable during later training on CIFAR-10.

This slower initial convergence is qualitatively similar to the learning behavior reported for BrainProp [Pozzi et al., 2020]. In BrainProp, learning is slower than conventional error backpropagation because only the output unit corresponding to the selected action is trained on a given trial, and the correct class is not revealed following an incorrect selection. REP shares these information constraints: conventional EP receives the complete target vector and therefore obtains an error signal for all output units, whereas REP receives only the selected class and a binary reward. However, the mechanisms used to propagate the resulting learning signal differ between the two approaches. BrainProp uses an explicit attention-gated feedback rule, whereas REP propagates the selected-output perturbation through the recurrent dynamics of symmetric three-phase EP. The restricted, action-dependent learning signal may therefore contribute to the slower initial convergence of REP, although further analysis would be required to determine the precise cause.

The generalization-gap analysis provided further insight into the differences between REP and EP. On MNIST and Fashion-MNIST, the two methods exhibited broadly similar gaps between training and test accuracy during the later stages of training. REP showed greater variability during some early and intermediate epochs, but the gaps of the two methods became comparable as training progressed. These results suggest that replacing the complete target vector with a selected-output reward signal did not substantially alter the final training–test discrepancy on these datasets.

A clearer difference was observed on CIFAR-10. EP reached a higher peak test accuracy during the earlier stages of training but showed a gradual decline after approximately 40 epochs. REP also exhibited a small decline after reaching its peak, but its test performance remained more stable during later training. In addition, REP consistently exhibited a smaller generalization gap than EP during the second half of training. By the end of training, the gap was approximately 28–29 percentage points for EP and approximately 23 percentage points for REP. This result is consistent with the comparatively stable test accuracy observed for REP during later training.

One possible explanation is that the distinction between REP and EP becomes more visible on CIFAR-10 because this dataset is more susceptible to overfitting than MNIST and Fashion-MNIST under the architectures and training settings used in the present study. CIFAR-10 contains higher-dimensional color images and greater within-class variability, while the convolutional architecture used for this task has sufficient capacity to fit the training data increasingly well over prolonged training. In such a setting, differences in the strength and specificity of the learning signal may have a greater effect on the relationship between training and test performance. Because REP updates the network using only the selected class and its associated binary reward, its restricted action-specific signal may limit the extent to which the model fits dataset-specific training patterns. This could produce a regularization-like effect and make the generalization-gap difference more apparent on CIFAR-10 than on the simpler MNIST and Fashion-MNIST tasks.

The smaller generalization gap of REP on CIFAR-10 should nevertheless be interpreted cautiously. A reduced gap does not necessarily imply improved generalization, because it may partly result from lower training accuracy. The present results do not establish whether the observed behavior arises from beneficial regularization, weaker optimization, reduced memorization, or another property of the reward-based perturbation. Additional analyses of training accuracy, training loss, model calibration, performance under different training durations, and behavior across architectures with different capacities would be needed to distinguish among these possibilities.

We also examined the effect of the exploration probability used by the Max–Boltzmann controller. The results showed that fully greedy class selection was not consistently optimal. Moderate exploration improved the mean performance on MNIST and Fashion-MNIST, while CIFAR-10 performance was relatively insensitive to the tested range of exploration probabilities. However, the relationship between exploration and performance was not monotonic, and some settings produced substantially larger variability across random seeds on Fashion-MNIST. These findings suggest that exploration can be beneficial, but its optimal value depends on the dataset and training conditions.

Although REP uses a reward-based learning signal, the present experiments do not constitute a complete reinforcement-learning setting. Each input is treated as an independent classification problem, the reward is delivered immediately after class selection, and there are no state transitions, delayed rewards, discount factors, or critic networks. REP should therefore be interpreted as a reward-based classification framework that provides an intermediate step toward reinforcement learning with EP.

An important future direction is to extend REP to sequential decision-making problems. Only a small number of studies have investigated the combination of EP and reinforcement learning [Kubo et al., 2022, Han and Sengupta, 2026]. REP may provide a useful foundation for this direction because it already replaces the complete supervised target with an action-specific reward signal. Future work could examine whether the same formulation can be extended to environments involving delayed rewards, multiple time steps, and temporal credit assignment.

Several biologically motivated extensions may also be considered. Dendritic neurons could be incorporated into REP by building on dendritic EP models [Kubo, 2026a]. Dendritic compartments may provide a mechanism for integrating feedforward, recurrent, and reward-related signals within individual neurons. Predictive learning rules could also be combined with REP to investigate whether local predictions can improve learning from sparse reward signals [Luczak et al., 2022, Luczak and Kubo, 2022, Kubo et al., 2023]. In addition, REP could be extended to wave-based recurrent neural networks (wRNNs), which provide recurrent dynamics for processing temporally structured information [Keller et al., 2024]. Such an extension may help bridge the present reward-based classification framework with sequential decision-making tasks. Dendritic wRNN architectures could further combine wave-like recurrent dynamics with compartmentalized neuronal processing, potentially enabling more flexible integration of temporal, contextual, and reward-related signals [Kubo, 2026b]. Heterogeneous neuronal time constants may also provide richer temporal dynamics and improve the stability of the free and nudged phases [Perez-Nieves et al., 2021, Kubo et al., 2026]. Finally, sleep replay consolidation could be explored as a mechanism for strengthening or reorganizing reward-associated representations after training [Tadros et al., 2022, Kubo et al., 2025].

These extensions should be evaluated carefully rather than being assumed to improve performance or biological plausibility automatically. In particular, future studies should determine whether each mechanism provides measurable benefits in learning efficiency, stability, generalization, or compatibility with local neural dynamics. Overall, the present results demonstrate that EP can learn from a selected-output binary reward signal while retaining performance reasonably close to conventional supervised EP. REP therefore provides a promising foundation for studying reward-based credit assignment in recurrent energy-based neural networks.

## Acknowledgments

This research was enabled in part by computational resources provided by the Digital Research Alliance of Canada (alliancecan.ca).

